# Flavin-containing siderophore-interacting protein of *Shewanella putrefaciens* DSM 9451 reveals substrate specificity in ferric-siderophore reduction

**DOI:** 10.1101/2024.05.13.594011

**Authors:** Inês B. Trindade, Bruno M. Fonseca, Teresa Catarino, Pedro M. Matias, Elin Moe, Ricardo O. Louro

## Abstract

*Shewanella* are bacteria widespread in marine and brackish water environments and emergent opportunistic pathogens. Their environmental versatility is highly dependent on the ability to produce an abundance of iron-rich proteins, mainly multiheme *c*-type cytochromes. Although iron plays a vital role in the ability of *Shewanella* species to survive in various environments, very few studies exist regarding the strategies by which these bacteria scavenge iron from the environment. Small molecule siderophore-mediated iron transport is a strategy commonly employed for iron acquisition, and it was identified amongst *Shewanella* spp. over two decades ago. *Shewanella* species produce hydroxamate-type siderophores and iron removal from these compounds can occur in the cytoplasm via Fe(III)-siderophore reduction mediated by siderophore-interacting proteins (SIPs). The genome of *Shewanella putrefaciens* DSM 9451 isolated from an infected child contains representatives of the two different cytosolic families of SIPs: the flavin-containing siderophore interacting protein family (SIP) and the iron−sulfur cluster-containing ferric siderophore reductase family (FSR).

Here, we report the expression and purification of the flavin-containing (*Sb*SIP) and iron-sulfur cluster-containing (*Sb*FSR) Fe(III)-siderophore reductases of *Shewanella putrefaciens* DSM 9451. The structural and functional characterization of *Sb*SIP shows distinct features from the highly homologous SIP from *Shewanella frigidimarina* (*Sf*SIP). These include significant structural differences, different binding affinities for NADH and NADPH, and lower rates of Fe(III)-siderophore reduction, results which consolidate in the putative identification of the binding pocket for these proteins.

Overall our work highlights NADH and NADPH specificity and the different Fe(III)- siderophore reduction abilities of the SIP family suggesting a tailoring of these enzymes towards meeting different microbial iron requirements.

## Introduction

Iron is one of the most abundant elements in the Earth’s crust, and its biogeochemical cycling mainly interconverts between the ferrous, Fe(II), and the ferric form, Fe(III). Iron is a key element for life, participating in various biological processes by mediating fundamental redox reactions via the incorporation into numerous proteins (1–3). Despite its abundance, in an oxygen-rich atmosphere like the present one, iron precipitates in its ferric form, and thus, it is not readily bioavailable (4, 5). To overcome iron shortage, microorganisms release siderophores, i.e., secondary metabolites with a high affinity for ferric iron which solubilize and scavenge iron for intracellular uptake (6–8). The main steps of the siderophore pathway include their intracellular synthesis, extracellular release into the environment, iron complexation, cellular uptake, and cellular iron release through specialized enzymes (9, 10). These can belong to the Siderophore-Interacting Protein (SIP) family or the Ferric-Siderophore Reductase family (FSR). So far, few studies exist regarding the function of FSRs and thus, the function of these two families of enzymes is considered redundant: to reduce ferric iron inside the Fe(III)-siderophore complex to its ferrous form, lowering the affinity of the complex and thus, promoting iron release (11–15).

The *Shewanella* genus is well known for its respiratory versatility, an ability that is facilitated by an extensive repertoire of iron-containing proteins, specifically, multiheme cytochromes (16, 17). Because of this, *Shewanellaceae* have higher iron requirements than many well-known bacteria, *e.g.,* approximately four-fold more than *Escherichia coli* (18). Despite their importance, there is still much to be learned regarding the iron acquisition pathways of this genus. Two decades ago, the iron-sequestering abilities of 51 strains of *Shewanella putrefaciens* isolated from different sources (fish, water, and warm-blooded animals) were assessed, where more than half of the strains produced hydroxamate-type siderophores (19). It is now becoming increasingly recognized that to thrive in a wide diversity of iron-deficient habitats ranging from the ocean bed to the eukaryotic host, *Shewanella* species adapt metabolically, including through the production of different siderophores and subsequent different proteins for their utilization (20, 21). Some examples include the production of cyclic dihydroxamate putrebactin by *S. putrefaciens*, and the production of asymmetrical avaroferrin by *S. algae*. Also, different *Shewanella* species have different Fe(III)-siderophore reductases (22). For instance, *S. frigidimarina* produces only a representative of the SIP family (*Sf*SIP), whereas some species produce representatives of both SIP and FSR families (11, 23–26).

In this work, we investigated siderophore-iron release from *S. putrefaciens* (DSM 9451) also known as *Shewanella* sp. JAB-1 and tentatively classified as *S. bicestrii* based on its genome sequence (27). *S. bicestrii* was first identified as part of three Extended Spectrum β-lactamase (ESBL)-producing bacteria in a bile sample of a 10-year-old child suffering from cholangitis (28).

Here we report the production of both Siderophore-Interacting Proteins, the flavin-containing (*Sb*SIP) and iron-sulfur cluster-containing Ferric-Siderophore Reductase (*Sb*FSR) from *S. biscestrii*. Given the instability of the latter, we only report the structure and biochemical characterization of *Sb*SIP. *Sb*SIP showed distinct binding constants for NADH and NADPH, and slower rates of Fe(III)- siderophore reduction when compared to previously characterized *Sf*SIP (14,15). Furthermore, the structure of *Sb*SIP revealed folding and electrostatic characteristics that together limit the access to the FAD cofactor which results in specificity towards NADH.

## Results

### Production of Fe(III)-siderophore reductases *Sb*SIP and *Sb*FSR

*Sb*SIP was heterologously expressed in *E. coli* and purified to apparent purity (>95%). It migrated as a single band at approximately 30 kDa (42 kDa before the HRV 3C protease incubation that removed the thioredoxin-His_10_ tag) on a 15 % SDS–PAGE gel, as expected from theoretical calculations (Fig. 1B). The purified protein appeared yellow and the UV–visible spectrum showed the typical spectral features of an oxidized flavoprotein in the UV–visible region (Fig. 1A), with absorption peaks at 387 nm and 471 nm and distinct shoulders located at 445 nm and 502 nm. *Sb*FSR was not purified to complete purity (Fig. 1D) given that the sample was very unstable and precipitated shortly after the first purification step. Regardless, a single band of protein migrated on a 15 % SDS– PAGE gel with the expected molecular weight of approximately 50 kDa (*Sb*FSR plus thioredoxin-His_10_ tag), as expected from theoretical calculations (Fig. 1D). The fractions containing *Sb*FSR were reddish brown, and the UV–visible spectra showed the typical spectral features of an oxidized 2Fe-2S protein, with maximum absorption peaks at 340 nm and 451 nm (Fig. 1C). The sequence of *Sb*FSR shows identities of 35 % to FhuF (*Escherichia coli* K-12) and 22 % to FchR (*Alkalihalophilus pseudofirmus*). It has an identical cluster binding motif sequence (C-C-x_10_-C-x_2_-C) and AlphaFold2, predicts overall folding conservation when compared to the archetypical FhuF from *E. coli* (29, 30). However, *Sb*FSR contains three extra α-helices and a longer N-terminal loop (Supplementary Information, Fig. S1) differences which may contribute to the observed protein instability.

**Fig. 1.**
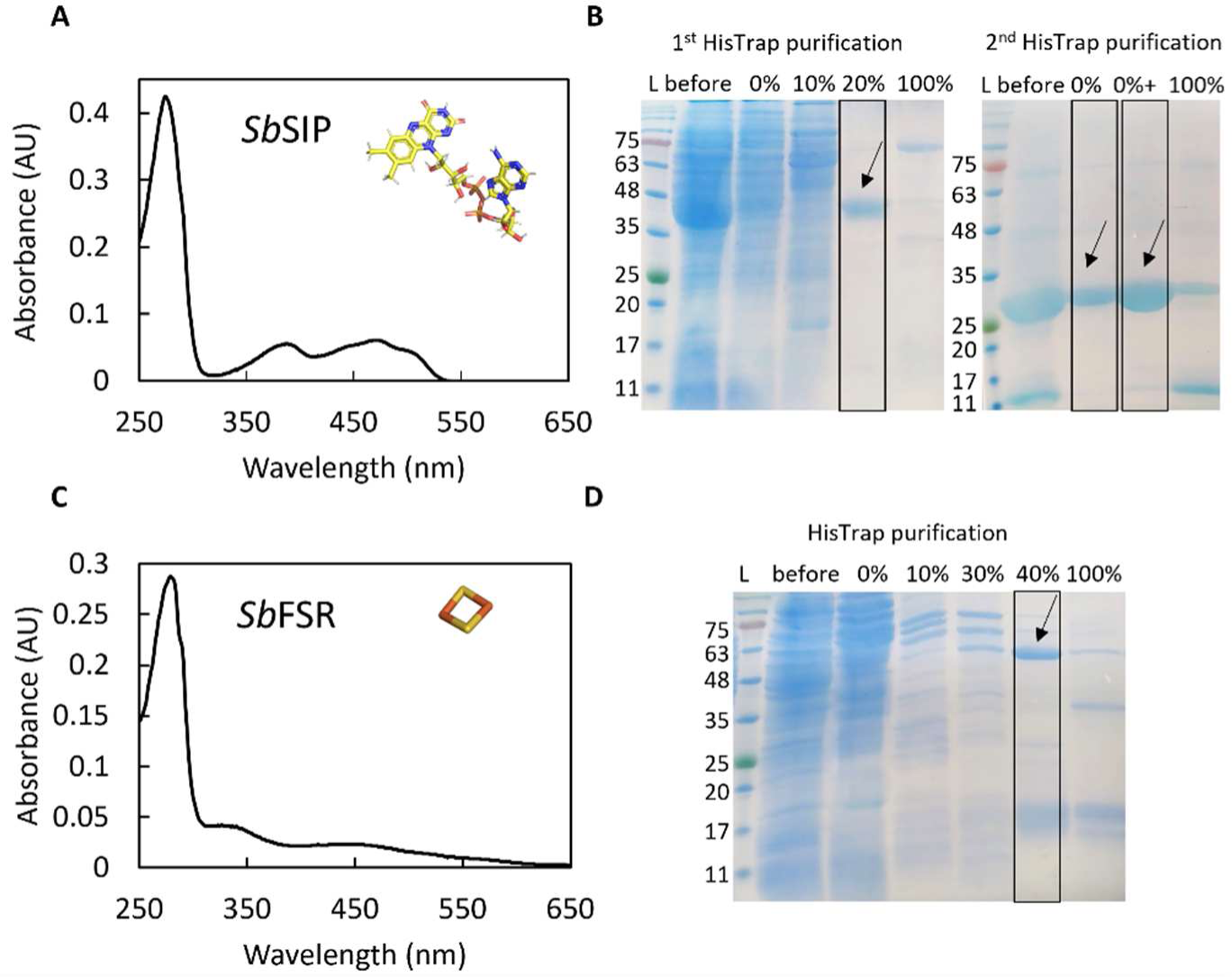
Production of *Sb*SIP and *Sb*FSR. **A)** UV-visible profile of pure *Sb*SIP showing the typical spectrum with maxima close to 387 nm and 471 nm. The insert depicts the stick representation of the spectroscopically active cofactor, the FAD **B)** SDS-PAGE gels of purification steps of *Sb*SIP before (1^st^ HisTrap purification) and after HRV 3C incubation step (2^nd^ HisTrap purification). **C)** UV-visible profile of the fraction containing *Sb*FSR showing the typical bands at 340 nm and 451 nm. The insert depicts the stick representation of the spectroscopically active cofactor, the 2Fe2S cluster **D)** SDS-PAGE gel of *Sb*FSR purification. Boxes represent fractions used for further experiments and arrows show gel bands of proteins of interest. Percentages represent the amount of imidazole used out of a 500 mM stock solution and the plus sign in panel B represents the sample after concentration.

### The structure of *Sb*SIP reveals a distinct electrostatic surface potential

The structure of *Sb*SIP was determined at 1.9 Å resolution (Fig. 2, Table 1). As previously described for other SIPs, the structure revealed two domains: N-terminal or NAD(P)H-binding and C-terminal or FAD-binding domain. The NAD(P)H-binding domain consists of the typical β1–α1–β2 Rossmann fold architecture composed of a manifold of β-antiparallel-strands (β1–β6) and two short α-helices (α1 and ƞ1). The FAD-binding domain is composed of five antiparallel β-strands (β7–β11) with two short α-helices (α2 and α3) connected by long loops.

**Fig. 2.**
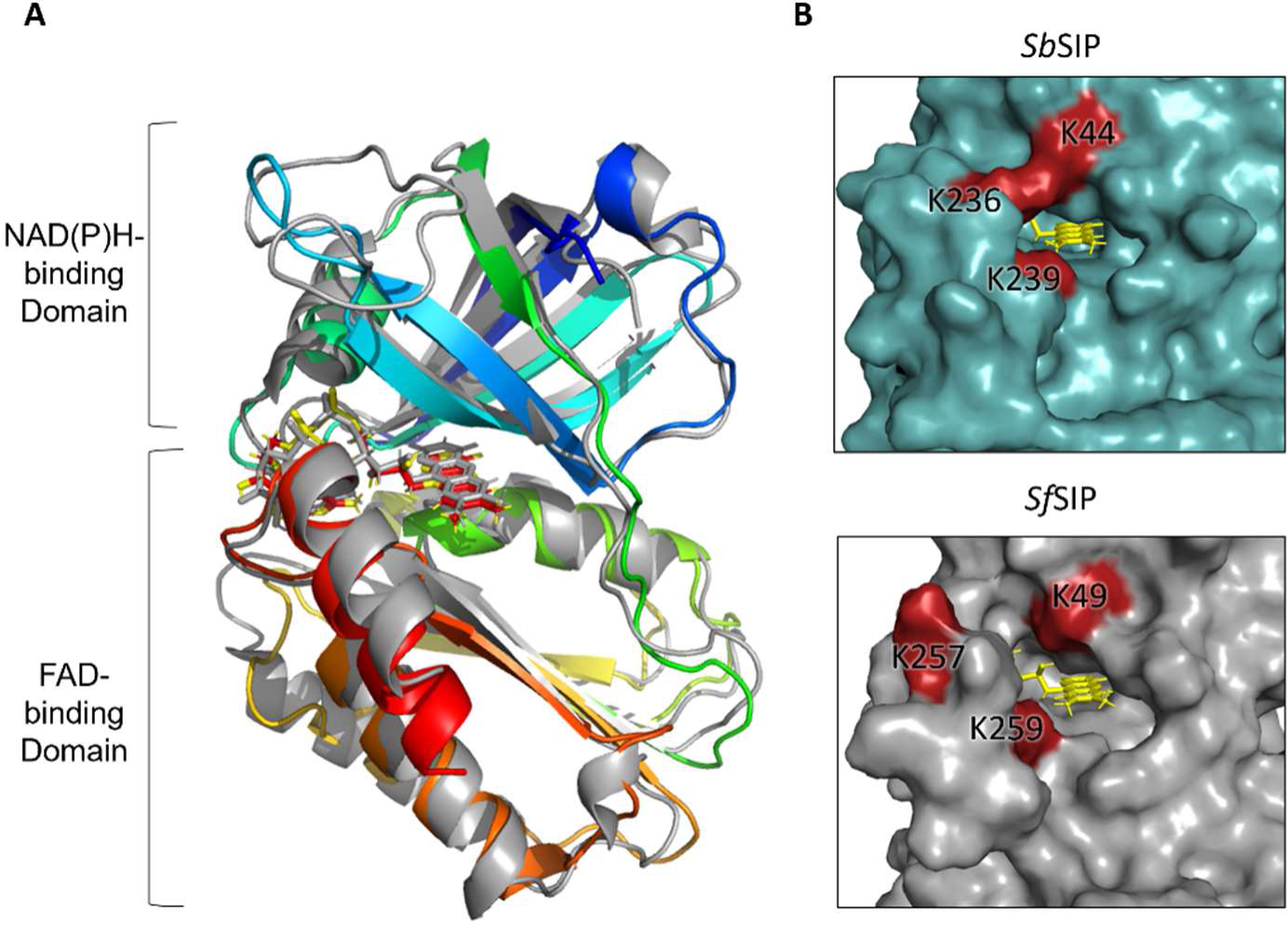
Structural characterization of *Sb*SIP. **A)** Structure of *Sb*SIP (blue to red from the N to the C terminal, PDB 8C4L) versus *Sf*SIP (gray, PDB 6GEH) aligned with PyMOL **B)** Molecular surfaces of *Sb*SIP and *Sf*SIP highlighting the lysine triad pockets (red) (31).

**Table 1.** Final refinement statistics of *Sb*SIP.

|  |  |
| --- | --- |
| Resolution limits (Å) | 62.71 – 1.86 (1.90 – 1.86) |
| R-factor (%) <sup>a</sup> | 16.9 (20.0) |
| nr. Reflections | 46650 (155) |
| Free R-factor (%) <sup>b</sup> | 17.4 (27.6) |
| nr. Reflections | 2412 (8) |
| Overall coordinate error estimate (Å) <sup>c</sup> | 0.17 |
| Model composition |  |
| Non-hydrogen protein atoms | 3826 |
| Solvent molecules | 323 |
| FAD ligand | 106 |
| Model r.m.s. deviations from ideality |  |
| Bond lengths (Å) | 0.012 |
| Bond angles (°) | 1.15 |
| Chiral centers (Å<sup>3</sup>) | 0.063 |
| Planar groups (Å) | 0.012 |
| Model completeness and validation |  |
| Regions omitted | 1-3 and 247-251 in chain A<br>1-5 and 249-251 in chain B |
| Mean B values (Å<sup>2</sup>) <sup>d</sup> |  |
| Protein | 39.0 |
| solvent molecules | 39.4 |
| FAD ligand | 28.6 |
| Ramachandran plot statistics. |  |
| Residues in: |  |
| most favored regions (%) | 96.3 |
| allowed regions (%) | 3.7 |
| disallowed regions (%) | 0.0 |
| Rotamer outliers (%) | 1.2 |
| C<sup>β</sup> outliers | 0.0 |
| Clash Score | 2.05 |
| Molprobity Score | 1.29 |

<sup>a</sup> R-factor = $\sum_{hkl} ||F_o| - |F_c|| / \sum_{hkl} |F_o|$ , where $|F_o|$ and $|F_c|$ are the observed and calculated structure factor amplitudes, respectively; <sup>b</sup> Free R-factor is the cross-validation R-factor computed from a randomly chosen subset of 5% of the total number of reflections, which were not used during the refinement.; <sup>c</sup> Maximum-likelihood estimate with PHENIX; <sup>d</sup> Calculated from the equivalent isotropic B values.

The FAD shows a planar conformation (Fig. S2), and it is stabilized through aromatic stacking interactions of the isoalloxazine ring with Tyr-60 and Tyr-225, and hydrogen bonds with residues Thr-61 (backbone O and N), Tyr-59 (side chain OH), Val-75 (backbone N), and Asp-73 (backbone O). The negatively charged phosphate groups in FAD are also stabilized through hydrogen bonds with Thr-60 (backbone N), Glu-232 (backbone N), His-77 (side chain Nε2), Gly-81 (backbone N), Ser-84 (Side chain OH), Ala-83 (backbone N). The triad of basic amino-acid residues (Lys-45, Lys-236, and Lys-239 in SIP from *S. putrefaciens, Sp*SIP) that has been proposed to form the Fe(III)-siderophore binding pocket is well-conserved (Lys-44, Lys-236 and Lys-239 in *Sb*SIP). As previously observed for *Sp*SIP, Lys-44 forms a ridge with Lys-236, making the access to the FAD cofactor smaller when compared to *Sf*SIP (Fig. 2B). Sequence alignment with previously characterized SIPs shows 82 % identity with *Sp*SIP (PDB code 2GPJ), 31 % with FscN (PDB code 4YHB), 30 % with *Sf*SIP (PDB code 6GEH), and 28 % with YqjH (no structure available). Indeed, when superimposing the structures of *Sb*SIP and *Sf*SIP, very few differences are observed in the overall fold. In contrast, the visualization of the electrostatic surface potential of these two proteins argues for remarkable differences (Fig. 3). Both *Sb*SIP and *Sf*SIP contain very positively charged pockets that provide access to the FAD cofactor through the isoalloxazine ring. However, the pocket of *Sb*SIP is substantially smaller when compared to *Sf*SIP. In *Sf*SIP it seems that the FAD cofactor is also accessible through the negatively charged phosphate groups and the pocket surface that surrounds it is still positively charged allowing NADH or NADPH the possibility of binding through both sides. In *Sb*SIP, access through the negatively charged phosphate groups seems more limited, and the electrostatic surface is negatively charged. It is likely that these differences in the access to the FAD cofactor are selection factors to discriminate redox partners, including Fe(III)- siderophores and the electron donors NADH and NADPH.

**Fig. 3.**
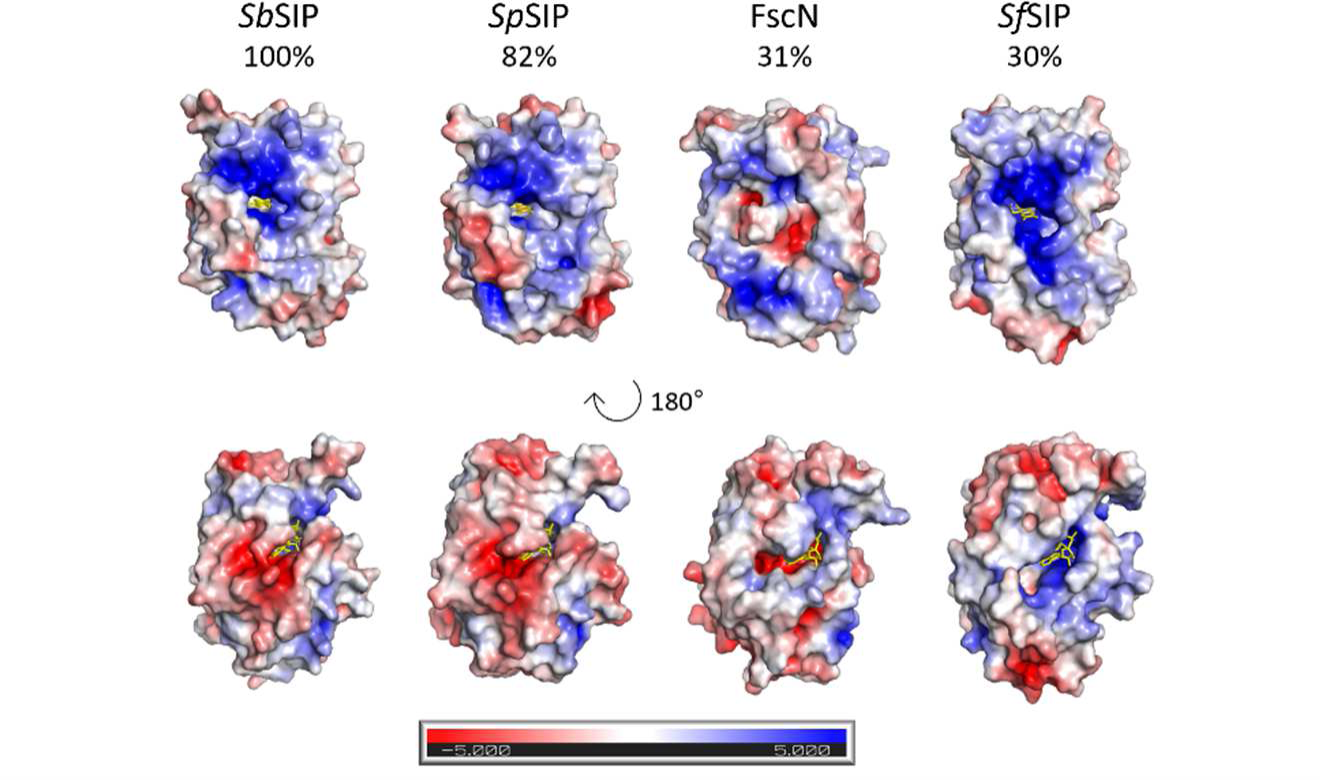
Electrostatic surface potential (−5 to +5 kT/e) of various SIPs, from *S. bicestrii* (*Sb*SIP, PDB 8C4L), *S. putrefaciens* (*Sp*SIP, PDB 2GPJ), *T. fusca* (FscN, PDB 4YHB) and *S. frigidimarina* (*Sf*SIP, PDB 6GEH). The surfaces are shown from most similar to least similar (left to right) in the same orientation. The top row highlights the access through the isoalloxazine ring, and the bottom row shows all surfaces after a clockwise 180 ° rotation about a vertical axis. Surfaces were calculated using the APBS plugin in PyMOL (32).

### *Sb*SIP performs proton-coupled electron transfer

Flavins are redox centers that can couple one- and two-electron transfer with proton transfer and this ability can be studied using Protein Film Voltammetry (PFV). *Sb*SIP presents two well-defined voltammetric signals (Fig. 4 A) and the midpoint potentials of these signals are reported in Table 2. This voltammetric response is significantly different from previously characterized *Sf*SIP where only one voltammetric signal was reported (24). Overall, from the reduction potentials obtained it is likely that one of the signals corresponds to the transition between the oxidized and the semiquinone state, and the other signal corresponds to the transition from the semiquinone to the hydroquinone state. The appearance of this latter signal that is absent in *Sf*SIP once again highlights the structural and electrostatic differences found in the surface of *Sb*SIP vs *Sf*SIP which can dictate interactions with the electrode surface and its access to the FAD cofactor. However, the remaining data showed comparable results to *Sf*SIP where one of the reduction potentials obtained is very similar (−210 mV vs SHE at pH 7 for *Sb*SIP and −228 mV vs SHE at pH 7, for *Sf*SIP, Table 2) and *Sb*SIP also presents a redox-Bohr effect extending throughout the physiological pH range, with predicted pK_ox_ lower than 4.5 and pK_red_ higher than 8.5 (Fig. 4B).

**Fig. 4.**
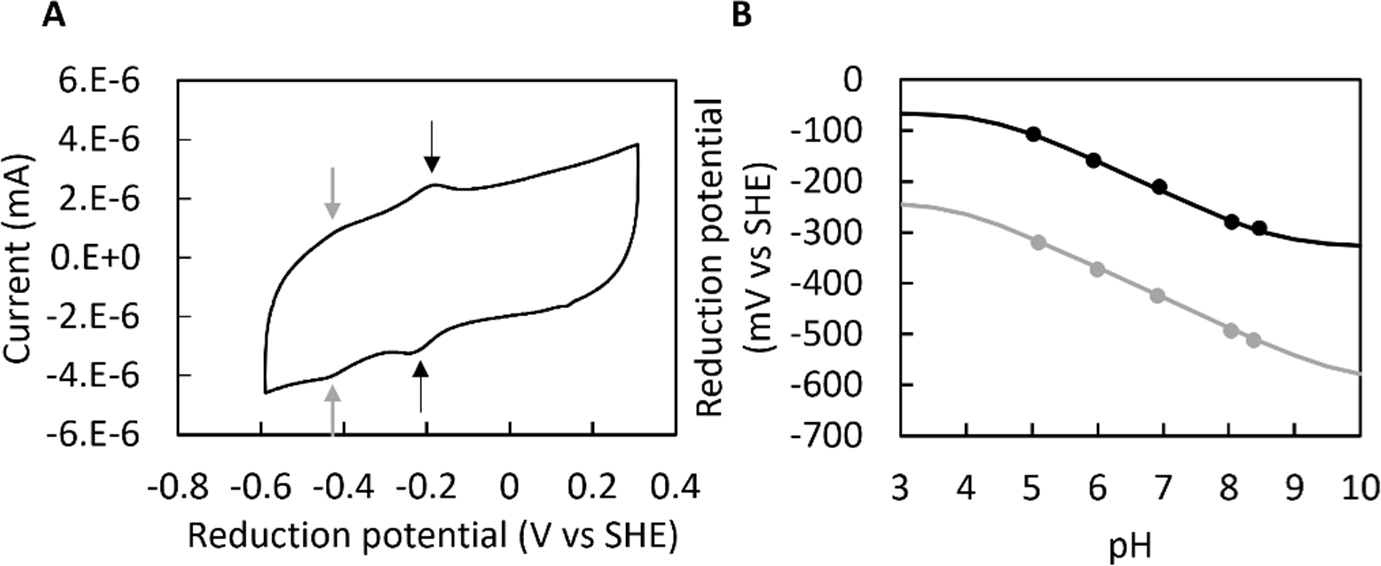
*Sb*SIP voltammetry experiments: **A)** representative voltammogram of *Sb*SIP, pH 7, at 100mV.s^-1^; gray and black arrows indicate the position of the low and high potential voltammetric signals, respectively, in the anodic and cathodic branch of the voltammogram. **B)** pH dependence of reduction potentials of *Sb*SIP with solid line representing the respective simulations, for the low potential signal (gray) and the high potential signal (black).

**Table 2.** pH dependence of the mid-point reduction potentials of *Sb*SIP. Values are averages of two experiments and respective standard error of the mean (SEM).

| pH | E1(1/2)(V vs SHE) | E2 (1/2)(V vs SHE) |
| --- | --- | --- |
| **5.02** | -0.106 ± 0.001 | -0.315 ± 0.005 |
| **5.94** | -0.163 ± 0.004 | -0.368 ± 0.005 |
| **6.94** | -0.214 ± 0.004 | -0.422 ± 0.003 |
| **8.04** | -0.280 ± 0.001 | -0.492 ± 0.002 |
| **8.46** | -0.294 ± 0.002 | -0.510 ± 0.003 |

### *Sb*SIP shows preference for NADH

*Sb*SIP interacts with NADH and NADPH in agreement with previously characterized SIPs, and binding to *Sb*SIP causes significant chemical shift changes in the ^31^P spectra of NADH and NADPH (Fig. 5 A and B). The binding of NADH to *Sb*SIP leads to changes in the ^31^P signals of the pyrophosphate in the slow-exchange regime in the NMR timescale, giving rise to the coexistence of resonances for the free and bound states. For NADPH the pyrophosphate signals and the 2’ signals displayed distinct behaviors, changing in slow-exchange regime for the former and in fast-exchange regime for the latter upon binding. The fact that the NADH and NADPH pyrophosphate ^31^P peaks change to similar positions upon binding to *Sb*SIP suggests a similar docking configuration of the two ligands in the bound state. The poor signal-to-noise of the NADPH spectra is a consequence of using lower concentrations of NADPH to achieve higher ratios of *Sb*SIP vs ligand which were necessary to characterize this weak binding event when compared with the binding of NADH. Consequently, the analysis of the chemical shift perturbation of the signal shifting in the fast-exchange regime showed greater accuracy and precision and was used to define the NADPH dissociation constant. The results indicate a nearly one-order-of-magnitude difference between the dissociation constants found for NADH and NADPH, of 17±5 µM and 107±2 µM, respectively. Both dissociation constants are consistent with transient interactions, although there is a clear preference of *Sb*SIP for NADH. This shows that *Sb*SIP is tailored towards receiving electrons that come predominantly from catabolic metabolism. This is in contrast with the previously characterized SIP from *S. frigidimarina* (*Sf*SIP) which did not discriminate between NADH and NADPH, with affinities of approximately 20 µM for both. Another contrasting feature is the fact that for *Sb*SIP the 2’ phosphate signal is perturbed upon binding, unlike what was observed for *Sf*SIP. The comparison between the FAD access pockets of *Sb*SIP and *Sf*SIP shows that the access to the FAD pocket in *Sb*SIP is smaller. Using molecular simulations, binding likely occurs near the isoalloxazine ring at distances sufficiently short to allow fast electron transfer, and both NADH and NADPH predictively bind in the same region (Fig. 6). Even though the distances between redox-active NADH, NADPH, and FAD cofactor are shorter for NADPH it is likely that the lower affinity for NADPH is a consequence of steric clashing. Furthermore, the binding conformations found can also explain the observed perturbation of the 2’ phosphate signal of NADPH when binding *Sb*SIP, since this extra phosphorus is facing towards the protein. In the case of *Sf*SIP the binding pockets found are similar to what was obtained for *Sb*SIP (Fig.S5) however in this case the 2’ phosphate of NADPH is facing outwards and therefore expected not to be disturbed upon binding (Fig. S6).

**Fig. 5.**
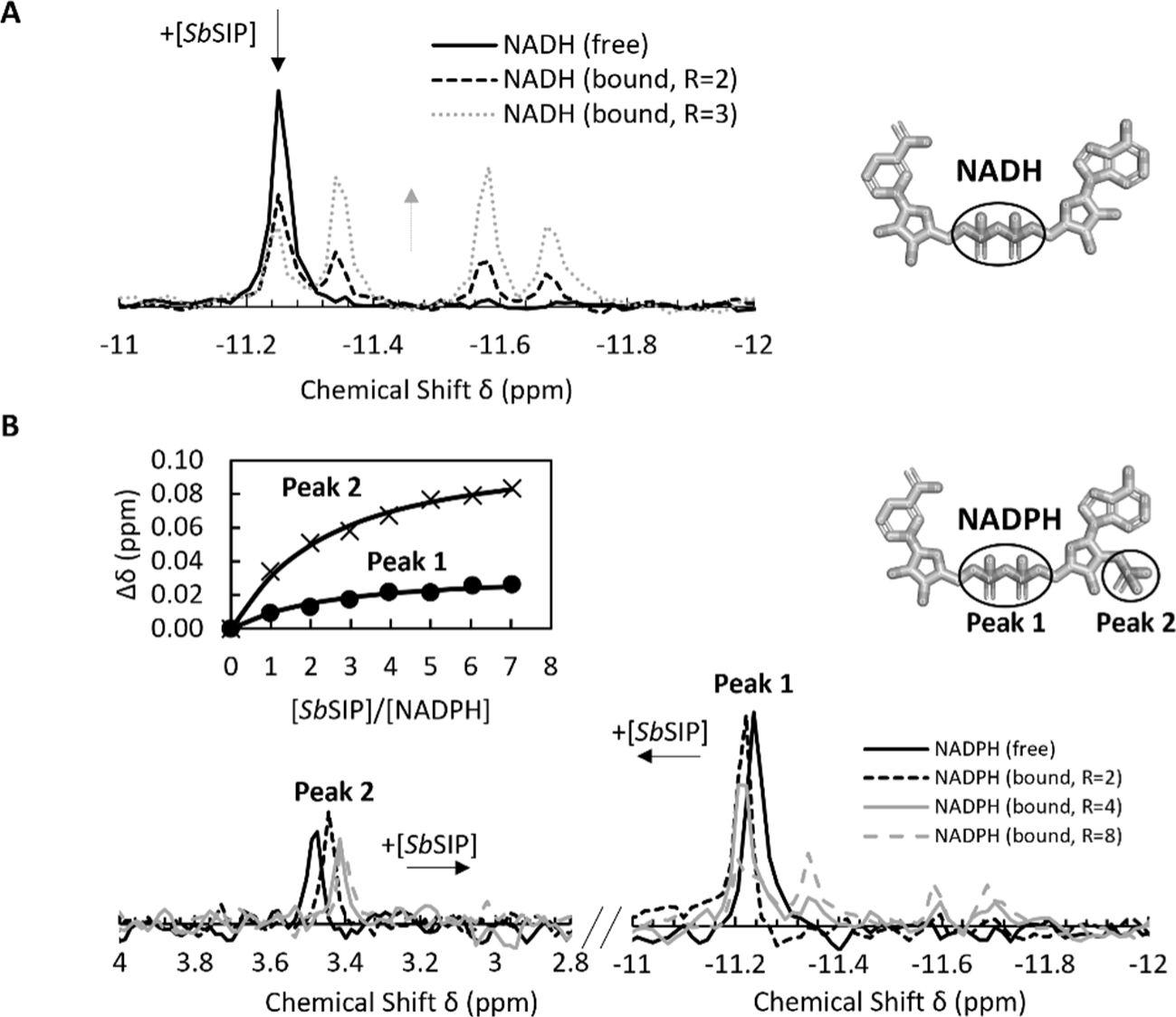
^31^P NMR binding experiments of NADH and NADPH versus *Sb*SIP. **A)** Proton-decoupled spectra corresponding to 100 μM of NADH and changes with increasing amounts of *Sb*SIP. R represents the ratio of [*Sb*SIP]/[NADH]. **B)** Proton-decoupled spectra corresponding to 57 μM of NADPH and changes with increasing amounts of *Sb*SIP. Insert shows binding curve monitoring the chemical shift perturbation. Peak 1 corresponds to the pyrophosphate phosphorous atoms, whereas peak 2 corresponds the 2’ phosphate group. R represents the ratio of [*Sb*SIP]/[NADPH].

**Fig. 6.**
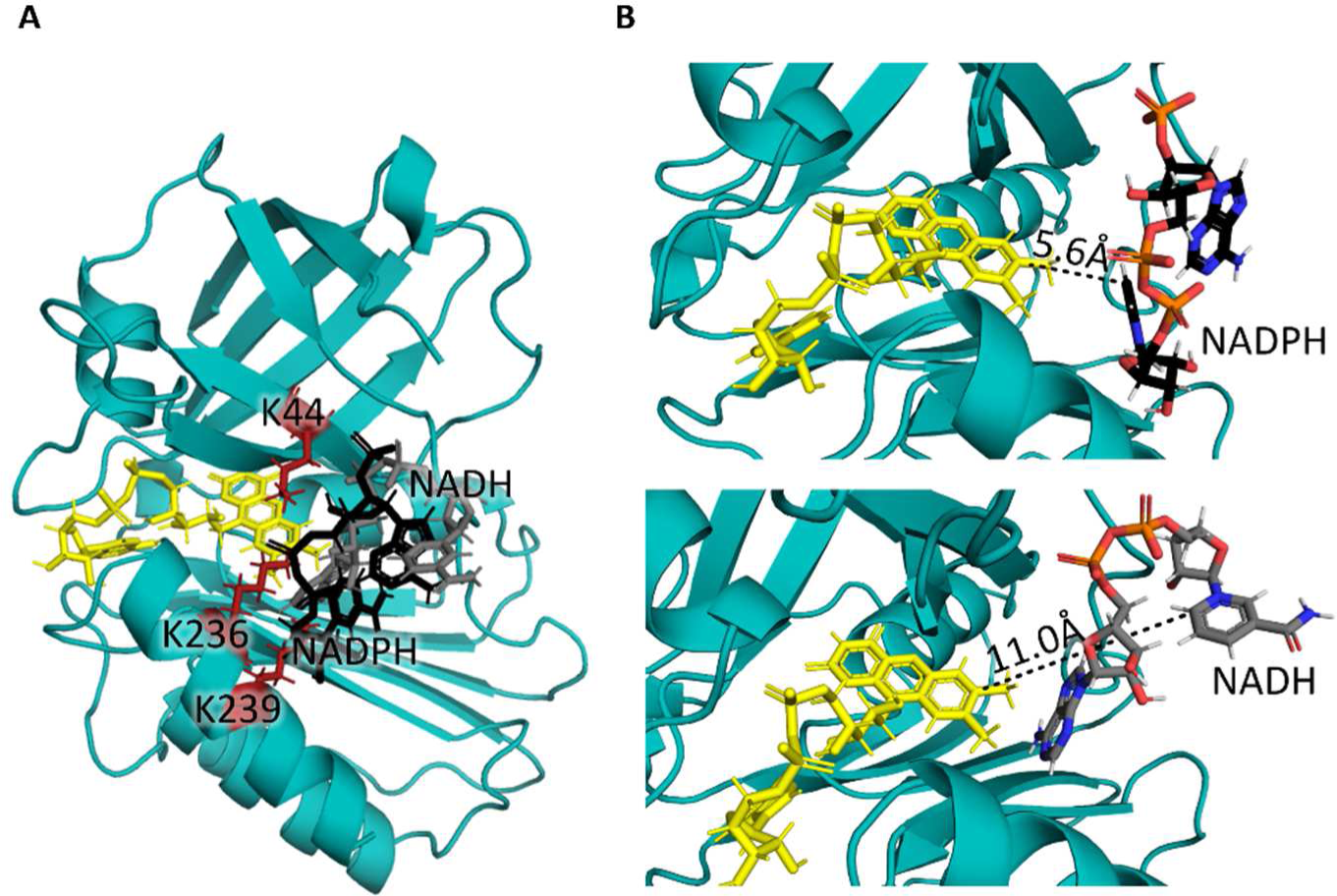
Representation of the binding conformations of NADPH and NADH to SbSIP calculated using Haddock. A) The binding of NADH (gray) and NADPH (black) occurs in the same region of the isoalloxazine ring of the FAD cofactor (yellow) which is surrounded by the lysine triad (red). (B) zoom of NADPH (top) and NADH (bottom) binding pockets highlighting the shortest distances found between the FAD cofactor and NADPH and NADH, respectively.

### SbSIP can be reduced by both NADH and NADPH

**Fig.6.**
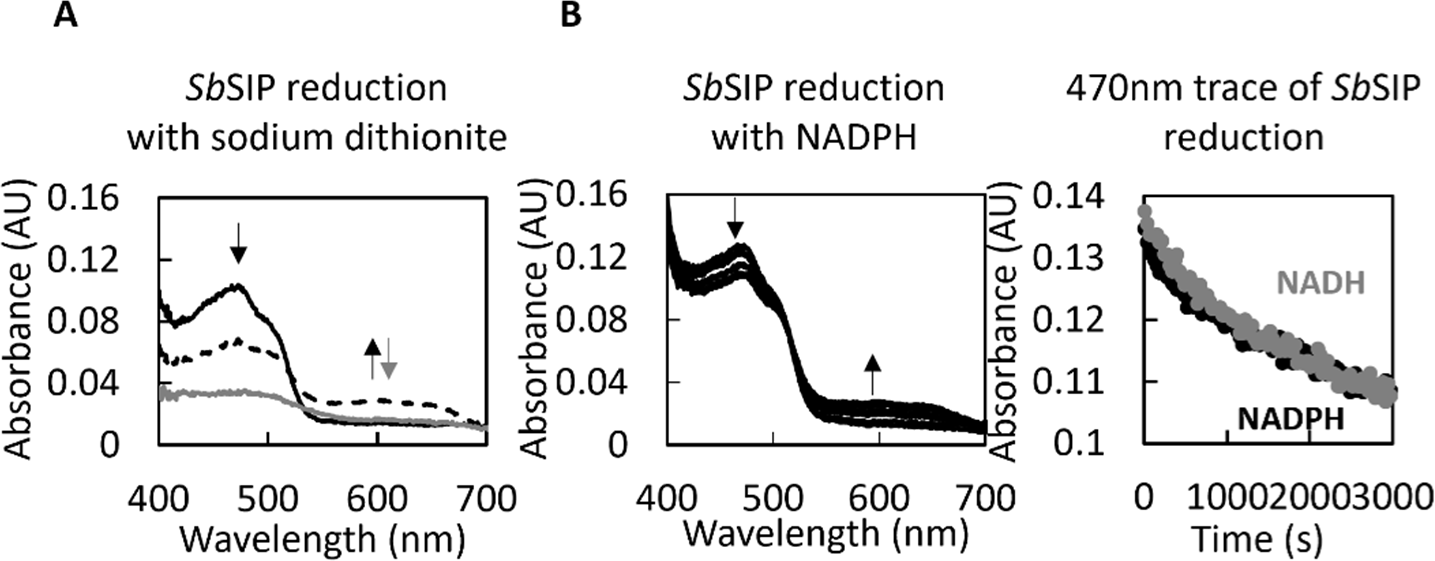
Reduction of *Sb*SIP with various electron donors. **A)** Spectral changes observed after mixing *Sb*SIP with sodium dithionite. The black line reports the spectrum of fully oxidized *Sb*SIP, which changes to a state with a significant amount of semiquinone population reported in the dashed line and then to the spectrum of the fully reduced state in gray. Black arrows indicate the decrease at 470 nm and increase at 600 nm occurring upon transition from fully oxidized to the semiquinone state, represented by the dashed line. Gray arrow indicates the decrease in absorbance both 470 nm and 600 nm upon transition from the semiquinone to the hydroquinone state, represented by the continuous gray line. **B)** Spectral changes observed after mixing *Sb*SIP with NADH and NADPH and respective kinetic traces at 470 nm. Data for NADH is reported in gray and for NADPH in black. Arrows indicate the direction of spectral changes upon reduction.

Given the differences found for the binding of *Sb*SIP with NADPH we then explored the reactivity of this enzyme. Upon mixing excess amounts of sodium dithionite with *Sb*SIP, absorption spectral changes were characterized by a decrease at 470 nm and an increase at 600 nm, followed by a decrease at both 470 nm and 600 nm. These changes are consistent with the full reduction of *Sb*SIP in two steps, the first is the formation of the semiquinone state followed by the formation of the hydroquinone state (Fig. 6A). The observation of the semiquinone state agrees with the significant separation between the two electrochemical signals observed in the PFV experiments (Fig. 4).

In the presence of the oxygen scavenging system, it was also possible to observe the reduction of *Sb*SIP with NADH and NADPH. Upon mixing excess amounts of NADH or NADPH with *Sb*SIP, absorption spectral changes were characterized by a decrease at 470 nm and an increase at 600 nm (Fig 6 and Fig. S3). These changes are consistent with the transition of *Sb*SIP from the oxidized to the semiquinone state. Full reduction into the hydroquinone state was not observed, in agreement with the more positive potential of NADH and NADPH vs the more negative voltammetric signal of *Sb*SIP (Fig. 4). Despite the differences in dissociation constant found in the ^31^P NMR binding experiments, no significant differences were found in the reduction rate of *Sb*SIP with NADH and NADPH. This likely occurs due to the use of excess reductant and therefore binding is not the rate-limiting step for *Sb*SIP reduction. The observation of *Sb*SIP reduction by NADH and by NADPH in the presence of the oxygen scavenging system led us to revisit the reactivity of the previously characterized *Sf*SIP which also showed the same behavior (Supplementary data Fig. S4). Interestingly, these results suggest that oxygen is an efficient inhibitor of SIPs. This provides a mechanism to avoid the deleterious effect of free ferrous iron in the cell in the presence of oxygen, i.e., the production of reactive oxygen species through the Fenton reaction, by preventing the formation of ferrous iron at the source.

### *Sb*SIP reduces bisucaberin and putrebactin slower than *Sf*SIP

The results of protein film voltammetry show that *Sb*SIP has the adequate thermodynamic properties to reduce ferric-siderophores. Therefore, its capability to perform the reaction was tested by stopped-flow kinetic assays. Upon mixing of *Sb*SIP_semi_ with ferric siderophores bisucaberin (BIS) and putrebactin (PUT) absorption spectra changes showed an increase at 470 nm and a decrease at 600 nm (Fig. 7A). This is consistent with the transfer of one electron from the semiquinone flavin to the Fe(III)-siderophore yielding fully oxidized *Sb*SIP and Fe(II). This result confirms the nature of *Sb*SIP as a NAD(P)H siderophore oxidoreductase. The rate constant for Fe(III)-siderophore reduction was determined to be 0.0014 ± 2×10^-4^ s^-1^ for BIS and 0.0031 ± 5×10^-5^ s^-1^ for PUT, from the kinetic trace at 600 nm to avoid spectral interference from the Fe(III)- siderophores (Fig. 7B). The reduction rate constants for both Fe(III)- siderophores are of similar magnitude, but are almost one order of magnitude lower than those found for *Sf*SIP (24). These results were surprising, since *Sb*SIP and *Sf*SIP display very similar reduction potentials. Given the similar driving force for the reaction, these results may reflect the structural differences between the two proteins. Despite its overall structural similarity to *Sf*SIP, surface access to the FAD cofactor is substantially diminished in *Sb*SIP (Fig. 3). Given that the rate of electron transfer decays exponentially with distance, an order of magnitude difference in electron transfer rates, as observed here, can arise from a small increase in distance, smaller than 3 Å, for the productive complex as described by the “Dutton-Moser ruler”(33). Indeed, these differences are observed using molecular dynamic simulations to predict the binding pocket of Fe(III)- siderophores in *Sb*SIP and *Sf*SIP. The binding conformations with the shortest distances between the isoalloxazine ring of *Sb*SIP and the Fe atoms in Fe(III)- bisucaberin are of 11.9 and 12.0 Å (Fig. 8) and 11.0 and 14.2 Å for the case of Fe(III)-alcaligin (Fig. S7), with both Fe(III)-siderophores binding to the same pocket. Contrarily, in the case of *Sf*SIP these distances are 9.0 and 12.5 for Fe(III)- bisucaberin and 9.0 and 13.0 Å for Fe(III)-alcaligin (Fig. S8 and S9), therefore comparatively shorter and consistent with faster electron transfer. It is also observed that in *Sb*SIP the positive cavity is smaller and less accommodating for one of the siderophore ligand moieties when in comparison with *Sf*SIP. Altogether, these differences suggest the tailoring of these proteins for different siderophore substrates (33, 34).

**Fig. 7.**
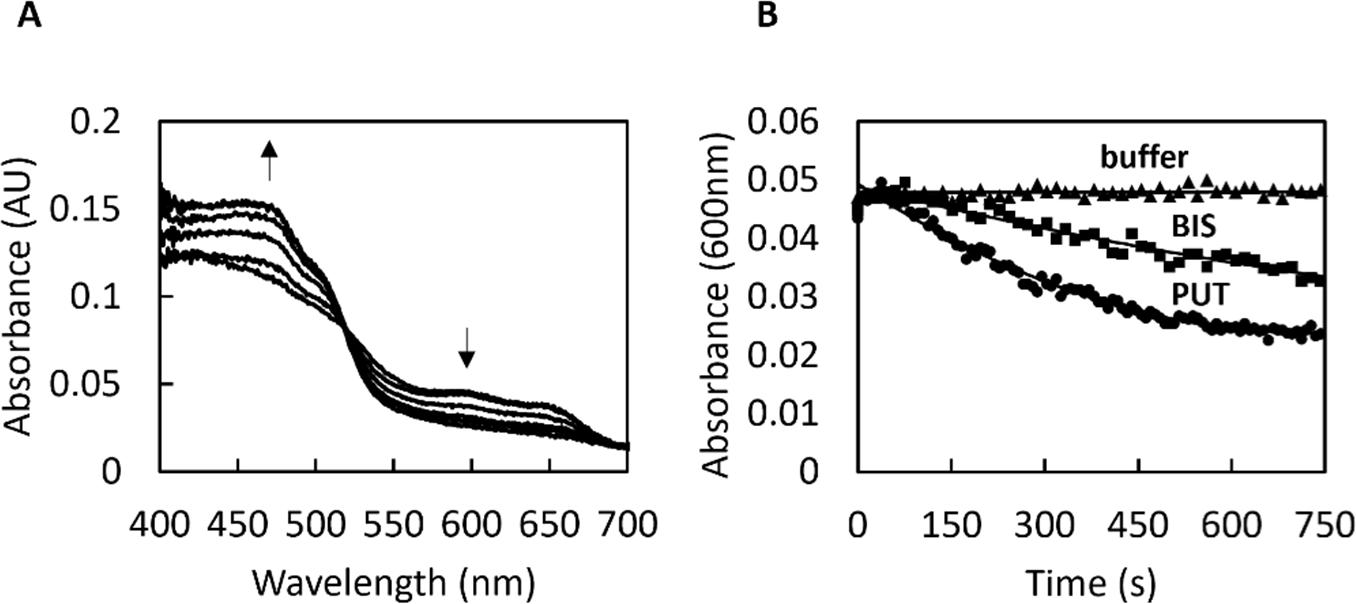
Fe(III)-siderophore reduction by *Sb*SIP: **A)** Absorption spectra changes after mixing *Sb*SIP_semi_ with Fe(III)-siderophore bisucaberin. Up arrow indicates the increase in absorbance at 470 nm and the down arrow indicates the decrease in the absorbance at 600 nm; **B)** Respective kinetic traces of *Sb*SIP at 600 nm, showing the oxidation of *Sb*SIP and concomitant reduction of bisucaberin (BIS) and putrebactin (PUT).

**Fig. 8.**
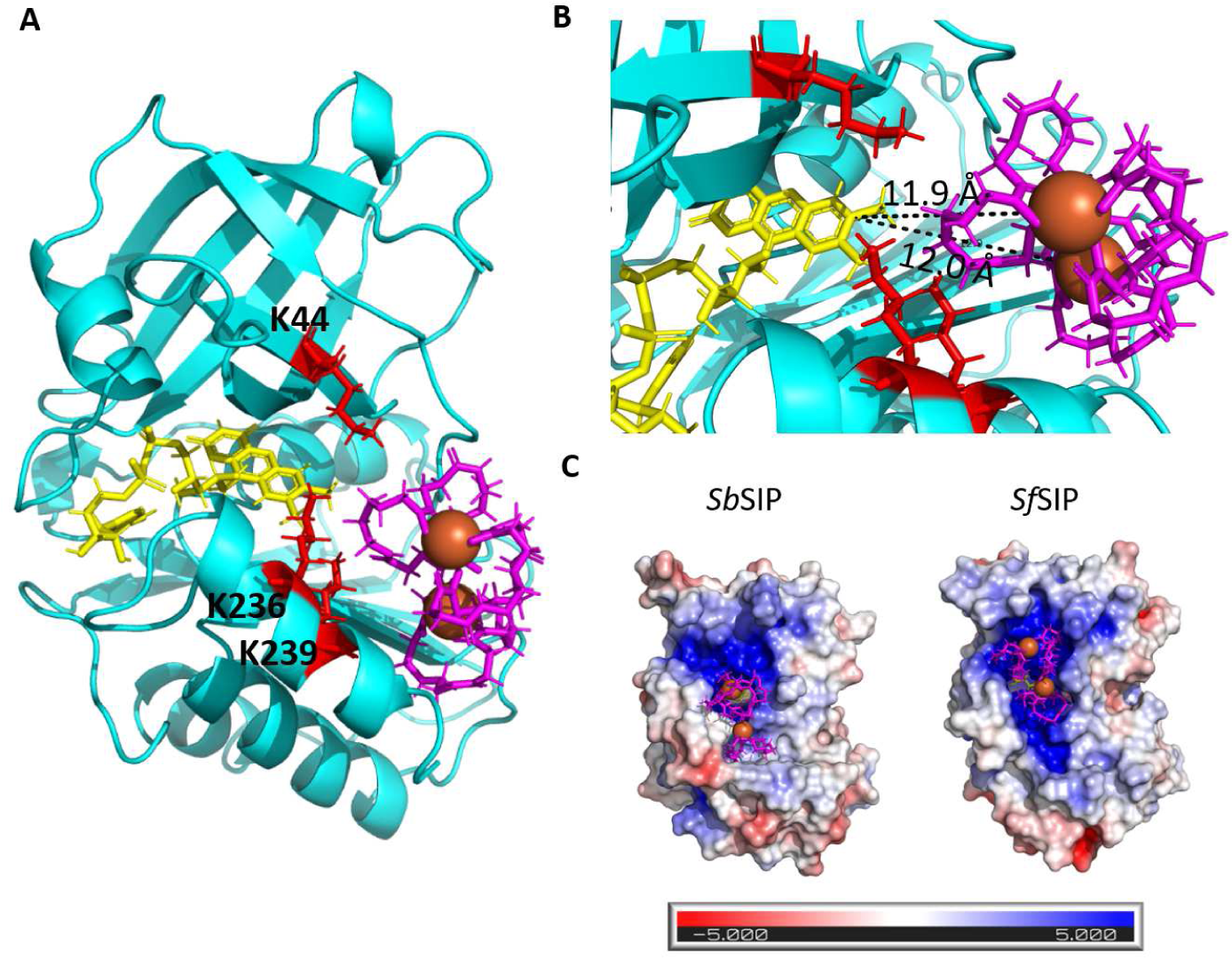
Representation of the binding conformations of Fe(III)-siderophores: **A)** Docking of Fe(III)-bisucaberin (pink) with SbSIP highlighting the lysine triad (red); **B)** Zoom of Fe(III)-bisucaberin binding pocket highlighting the shortest distances between the isoalloxazine ring of the FAD cofactor and the two Fe(III) atoms of the Fe(III)- bisucaberin complex; **C)** Electrostatic surface potential (−5 to +5 kT/e) of *Sb*SIP and *Sf*SIP and respective Fe(III)-bisucaberin binding pockets.

## Discussion

Here we present the functional characterization and structure determination of a novel SIP from *S. bicestrii* (*Sb*SIP). The comparison with the previously characterized SIPs highlights features that appear relevant for the functional mechanism and physiological role of these enzymes.

The observation of redox-Bohr effect as previously reported for *Sf*SIP suggests that this is a common functional feature of these enzymes. The coupled transfer of electrons and protons in the physiological pH range is clearly advantageous for the activity of this enzyme. The redox-Bohr effect implies that the driving force for Fe(III)-siderophore reduction is enhanced at high pH, where the solubility of Fe(III) is low. Moreover, it ensures that iron reduction is accompanied by the release of a proton resulting in local pH reduction and enhanced Fe(II) solubility.

The structure of *Sb*SIP presented an overall similar fold, but detailed structural differences were found when compared to previously characterized SIPs and this can be correlated with functional differences. A smaller FAD pocket likely sets the preference for NADH via steric clash with the 2’ phosphate group of NADPH and possibly a direct link to catabolism and/or preferential use of specific siderophores. Furthermore, an affinity for NADH like that found for *Sf*SIP reveals that it is through this positively charged pocket that binding takes place. Miethke and co-workers have previously suggested two different SIP subgroups based on the presence (subgroup I) or absence (subgroup II) of a longer C-terminal α-helical element (14). Subgroup I would favor NADH binding, and subgroup II would favor NADPH binding. Both SIP from *E. coli* (YqjH) and SIP from *T. fusca* (FscN) were shown to meet this criterion, belonging to subgroup II and binding NADPH-only. However, *Sf*SIP (SIP from *S. frigidimarina*) was shown to not meet the criterion, binding both NADH and NADPH with similar affinity, despite its classification as subgroup I (15). Interestingly, *Sb*SIP also belongs to subgroup I as well, but it shows a preference for NADH, suggesting that the extra C-terminal helix indeed hinders the access to the FAD pocket through the isoalloxazine ring. However, other factors play a role in tailoring the activity of SIPs towards NAD(P)H. This modulation of the affinities may stem from different poises in the NAD^+^/NADH and NADP^+^/NADPH ratios in different organisms and the regulation of reducing power flux through the catabolic or anabolic pathways according to their needs.

The siderophores used in this study are produced by members of the genus *Shewanella.* However, very few studies exist regarding the exact siderophores, and corresponding Fe(III)-siderophore structures produced by the different species of *Shewanella*, or even which microorganisms form symbiotic relationships with the different *Shewanella* species. Thus, the slow kinetics of siderophore reduction observed for SbSIP may reveal an alternative scenario to targeting the siderophores produced by the organism from which each SIP was isolated. SIPs may be also tailored to specifically target xenosiderophores as observed in the case of *E. coli*’s native siderophore enterobactin. Most microorganisms do not synthesize this siderophore but nonetheless express the ferric enterobactin esterase (Fes). This is also true for some *Shewanella* species, and it could provide a means to release iron when it is complexed with enterobactin, the strongest siderophore described to date (35).

Finally, it is also important to reflect upon the environmental context of different *Shewanella* species including where they were isolated from and their different iron requirements. *S. frigidimarina* was isolated from the Antarctic soil bed at very cold temperatures whereas *S. bicestrii* was isolated from the warmth of the human body (28, 36). The lower access to the FAD in *Sb*SIP may have been an evolutionary adaptation to slow down the rates of siderophore reduction at the higher temperature of the *S. bicestrii* habitat. This provides a kinetic control on the production of Fe(II) that may be essential to enable the metabolism to handle this essential but potentially toxic element while maintaining a similar thermodynamic driving force. Additionally, the lower reduction rates can also be a consequence of the fact that the genome of *S. bicestrii* codes for two ferric-siderophore reductases, *Sb*SIP and *Sb*FSR, whereas the genome of *S. frigidimarina* only codes for *Sf*SIP. Again, kinetic control of the relative activities of the two enzymes may be important in the metabolic regulation of iron availability, which operates on a much faster response time scale than transcriptional regulation. Nonetheless, it is also possible that the binding of siderophores is affected by temperature, either for their affinity to *Sb*SIP or due to the temperature dependence of speciation equilibria which changes iron availability.

Overall, this study reveals the specificity of SIPs for NADH and NADPH and how oxygen can be a strong inhibitor of the activity of these enzymes. It also highlights how the physicochemical context of iron availability together with the availability of different siderophores produced by the surrounding ecological community constitute key factors shaping the function of these enzymes. These factors are now ripe for exploration and will likely be the make-it-or-break-it in the context of using these enzymes as targets to combat bacterial infections and antimicrobial resistance.

## Experimental procedures

### Production of *Sb*SIP and *Sb*FSR

*Sb*FSR and *Sb*SIP expression vectors were designed based on the SLIC (Sequence and Ligation Independent Cloning) method developed by Scholz and co-workers (37). Primers were designed as described therein to include an HRV 3C protease cleavage site and ccdB counter-selection (3C-LP1 and ccdB-LP2). Gene fragments *sbsip* and *sbfsr* were amplified using KAPA2G Robust PCR kit from a colony of commercially available *S. putrefaciens* DSM 9451 (DSMZ) grown on a Luria Bertani (LB) agar plate. PCR fragments were then ligated into pCoofy38, a vector containing an N-terminal Thioredoxin-His_10_ tag using a Gibson Assembly® Cloning Kit (New England BioLabs). Plasmids were extracted and transformed into competent *E. coli* BL21(DE3) cells for expression. Proteins were expressed by growing expression strains in LB Broth Media (*Sb*SIP) or Terrific Broth Media (*Sb*FSR) supplemented with 50 mg/L kanamycin at 37 °C, 150 rpm, and expression was induced with 1 mM IPTG at an OD_600_ of 0.5-0.7. After approximately 30 hours, cells were harvested by centrifugation and frozen at - 80 °C. Cells were later defrosted and resuspended in 20 mM Potassium Phosphate buffer pH 7.6 with 300 mM NaCl and a protease inhibitor cocktail (Roche) together with DNase I (Sigma) prior to a three-pass cell disruption at 6.9 MPa using a French press. The lysate was ultracentrifuged at 204 709 x g for 75 min at 4 °C to remove cell membranes and debris. Proteins were purified from the supernatant using a His-trap affinity column (GE Healthcare) with a stepwise elution method. The fraction containing *Sb*FSR was eluted at 20 mM Potassium Phosphate pH 7.6, 300 mM NaCl with 250 mM imidazole whereas *Sb*SIP was eluted with 150 mM imidazole. Eluted fractions were analyzed by SDS-PAGE using Blue-Safe staining (NZYTech) and UV–visible spectroscopy to select fractions containing *Sb*FSR and *Sb*SIP. The fractions of each protein were pooled, the imidazole was removed through dialysis overnight, and proteins were concentrated using an Amicon ® Ultra Centrifugal Filter (Millipore) with a 30 kDa cutoff. The *Sb*FSR yield was very low, and fractions precipitated after concentration. Thus, Trx-His_10_ tag cleavage was only performed for *Sb*SIP. *Sb*SIP fractions were incubated with HRV 3C Protease overnight at 4 °C with agitation, and the final purified *Sb*SIP was concentrated from the flow-through of a second passage through the His-trap column using an Amicon Ultra Centrifugal Filter (Millipore) with a 30 kDa cutoff. The purity of *Sb*SIP was confirmed by SDS-PAGE using Blue Safe staining (NzyTech), N-terminal sequencing analysis, and UV-Visible spectroscopy. An extinction coefficient of free FAD ε_450nm_= 11 300 M^-1^ cm^-1^ was used for quantification purposes (38).

### Crystallization and structure Determination of *Sb*SIP

Purified *Sb*SIP (Trx-His10 tag-free) at a concentration of 10 mg/ml was crystallized by the hanging drop vapor diffusion technique using as precipitant a solution containing 1.8 M Ammonium sulfate with 0.01 M Cobalt(II) chloride hexahydrate and 0.1 M MES pH 6.5. Drops containing 1 μl protein and 1 μl reservoir were equilibrated against 500 μL reservoirs in a 24-well plate (Hampton Research). Crystals were harvested and soaked in a 2 μl drop of cryo solution (1.8 M Ammonium sulfate 0.1 MES pH 6.5 with 30 % glycerol) prior to flash-freezing in liquid nitrogen. Diffraction data were collected at 100 K to a resolution of 1.86 Å at the ALBA beamline XALOC (Barcelona, Spain). The images were processed with AutoProc and STARANISO, which make use of XDS and the CCP4 suite for integration and conversion of integrated intensities to structure factors (39–44). The data processing statistics are listed in Table 3. The structure was solved by molecular replacement using PHASER in the CCP4 suite and a previously determined SIP crystal structure from *S. putrefaciens* (PDB 2GPJ, Joint Center for Structural Genomics) as phasing model. The asymmetric unit cell of the crystal contained two *Sb*SIP protein chains (A and B), each one bound to a FAD moiety, and the model was automatically corrected with BUCCANEER/REFMAC in the CCP4 suite (45, 46).

**Table 3.**
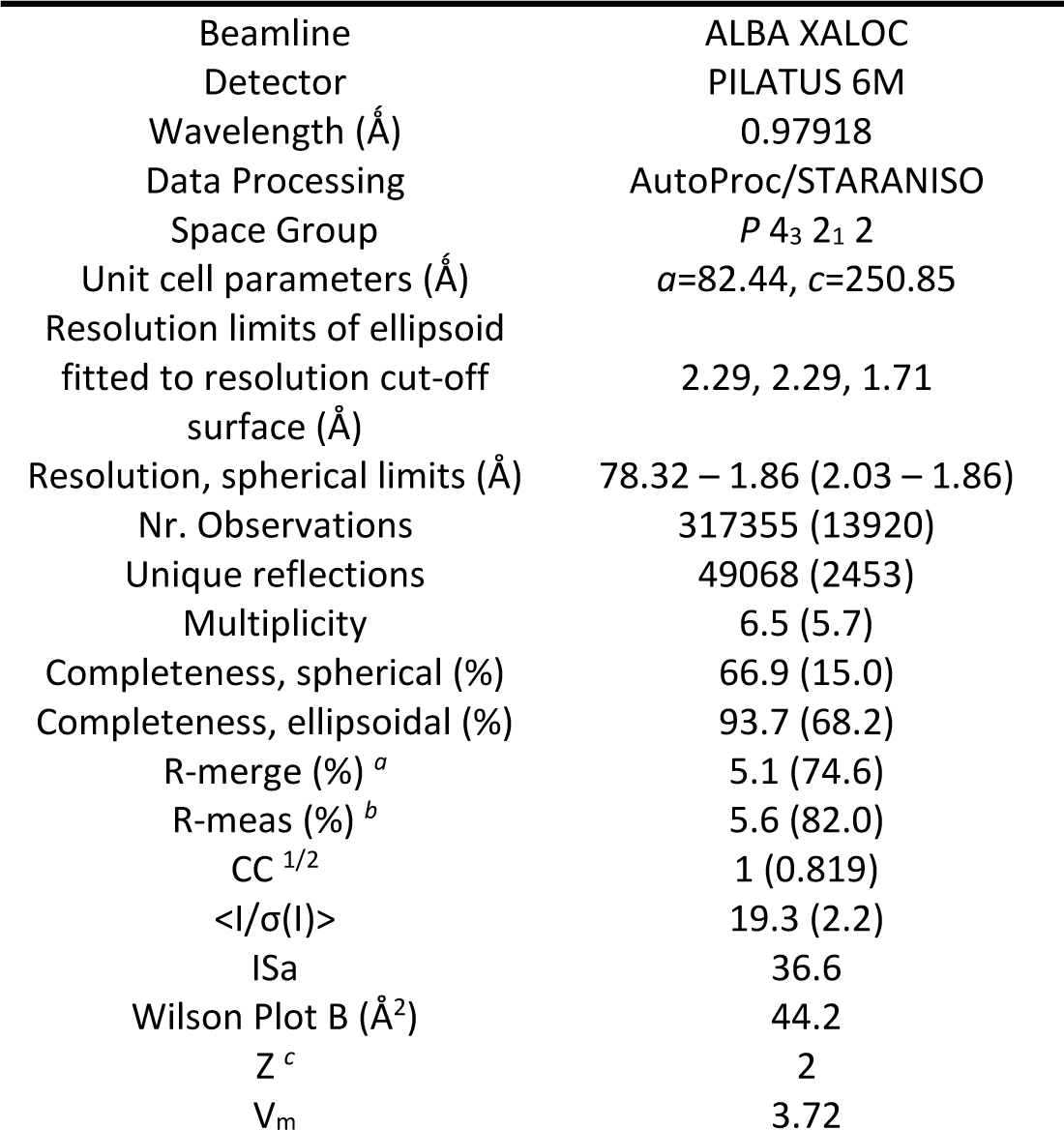
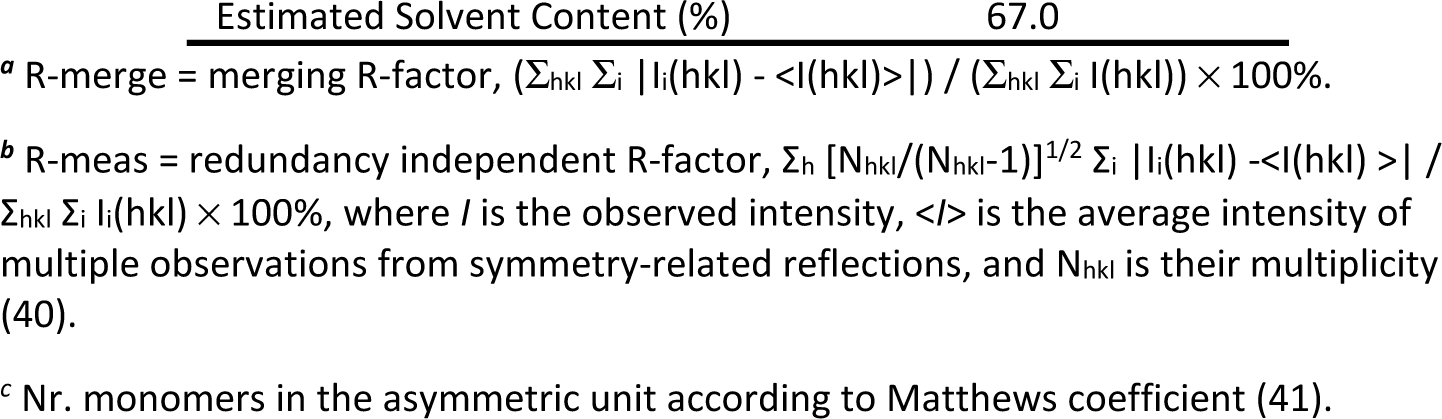
Data collection and processing statistics.

After an initial refinement using REFMAC5 in the CCP4 suite, structure refinement was continued using PHENIX (47). Hydrogen atoms were included in calculated positions with the PHENIX READYSET tool, and isotropic atomic displacement parameters (ADPs) were refined for all non-hydrogen atoms. Throughout the refinement, the model was periodically checked and corrected with COOT against σ_A_-weighted 2|F_o_|-|F_c_| and |F_o_|-|F_c_| electron-density maps. Solvent molecules were added automatically by the ArpWarp solvent protocol via the CCP4 suite and validated by inspection of electron-density maps in COOT (48, 49) In the final refinement cycles, a TLS rigid body refinement of the ADPs was carried out, considering 7 and 6 rigid body groups for *Sb*SIP chains A and B, respectively, determined with the PHENIX FIND_TLS_GROUPS tool from a previous refinement with isotropic ADPs. The final values of R and R-free were 0.168 and 0.200 respectively, with a maximum likelihood estimate of the overall coordinate error of 0.17 Å (50). The refinement statistics are presented in Table 3. The model stereochemical quality was analyzed with MOLPROBITY and there are no outliers in the Ramachandran ϕ,φ plot (51). The coordinates and structure factors have been submitted to the Worldwide Protein Data Bank (wwPDB consortium, 2018) with accession code 8C4L. Images were produced using PyMOL (52).

### Protein Film Voltammetry (PFV) of *Sb*SIP

PFV experiments of *Sb*SIP were performed at 25 °C using a three-electrode electrochemical cell configuration with a PGE (pyrolytic graphite edge) electrode, a graphite rod (counter electrode) and an Ag/AgCl 3 M KCl (reference electrode) inside a Coy anaerobic glovebox chamber using a CHI electrochemical analyzer (CHI instruments). The electrode was cleaned and freshly polished before every experiment. The polishing routine consisted of a 10 min nitric acid incubation at room temperature followed by 10 min of hand-polishing with a 1.0 μM alumina aqueous slurry. The electrode was thoroughly rinsed with water and left to dry and then *Sb*SIP was immobilized by pipetting 7 μL of a 250 μM solution of *Sb*SIP in 20 mM Potassium Phosphate buffer at pH 7.6 with 100 mM KCl. Once fully dried, the electrode was rinsed to remove protein excess and immersed in Potassium Phosphate buffer at different pH values. Experiments were performed at different scan rates and then the buffer was collected, and the pH was measured for confirmation. QSoas was used to subtract the capacitive current and extract the reduction potentials. Potentials are reported in mV versus the Standard Hydrogen Electrode (SHE) by the addition of 210 mV to those measured (53).

### 31P NMR: NAD(P)H binding experiments

NADH, NADPH and *Sb*SIP were prepared in 20 mM Tris-HCl buffer at pH 8 with 100 mM KCl containing 10 % of ^2^H_2_O (99.9 atom %). Using the standard Bruker pulse program “zgdc,” one-dimensional proton-decoupled ^31^P spectra were acquired with 2048 scans, d1 of 1.3 s at 25 °C on a Bruker Avance II 500 MHz equipped with a SEX probe for ^31^P detection. Samples of 100 µM or 57 µM of NADH and NADPH, respectively, were titrated against increasing concentrations of *Sb*SIP. Collected spectra were visualized and analyzed using TopSpin 3.6 (Bruker). For NADH binding, the concentration of free and bound species was determined from the relative intensity of each peak and the dissociation constant (K_d_) was calculated using the equation previously described (24). For NADPH binding, the chemical shift perturbations (Δδ) of the NMR signals from NADPH that resulted from the complex formation with *Sb*SIP in the fast exchange regime were plotted against the molar ratio (R) of [*Sb*SIP]/[NADPH]. Results were fitted and the dissociation constant and respective uncertainty (K_d_) was determined as described by Fonseca and co-workers (54).

### Kinetic Experiments

The kinetic experiments were performed with HI-TECH Scientific Stopped-flow equipment (SF-61DX2) installed inside an anaerobic glove box (Mbraun MB150-GI). The temperature of the drive syringes and mixing chamber was maintained at 25 °C using a water bath. Sample solutions were prepared with 20 mM Potassium Phosphate buffer pH 7 with 100 mM KCl and in the presence of an O_2_ scavenging system (10 mM glucose, 375 nM glucose oxidase, and 750 nM catalase). The time course of the reactions was monitored using a photodiode array. Solutions were prepared inside the anaerobic chamber with degassed water and all experiments were performed in triplicate. Data were analyzed with Kinetics Studio version 2.32 (TgK Scientific).

Reduction of both *Sb*SIP and *Sf*SIP with NADH and NADPH was performed by mixing 1 mM of these compounds with 20 μM of *Sb*SIP or *Sf*SIP. The reduction with sodium dithionite was performed by mixing 3 mM of this compound with 20 μM *Sb*SIP.

Fe(III)-siderophores putrebactin and bisucaberin were kindly provided by Prof. Masaki Fujita. Reduction of these by reduced *Sb*SIP (*Sb*SIP_semi_) was performed after reducing *Sb*SIP with sodium dithionite. The latter was achieved by using excess sodium dithionite which was then removed through buffer exchange in a HiTrap® Desalting Column (GE Healthcare). Ferric-siderophore reduction experiments were then performed using 20 μM *Sb*SIP_semi_ against 100 μM of ferric-siderophore in the stopped-flow apparatus. The reduction rate constants were obtained from the fitting of the kinetic traces at 600 nm. Catalytic experiments with NADH or NADPH and ferric-siderophores were attempted. However, spectra overlap prevented an unambiguous interpretation of the results, and thus, these data were excluded from this manuscript. Ferrozine assays were also attempted. However, in the presence of the oxygen-scavenging system, absorption changes were observed in the absence of *Sb*SIP (control). Ferrozine is a very strong ferrous iron chelator and in the absence of oxygen, the shift in chemical equilibrium is sufficient for Fe(III)-siderophore reduction to take place at detectable rates without enzymatic mediation (55).

### Docking NADH, NADPH and Fe(III)-siderophores

The ligand docking calculations were executed through the High Ambiguity Driven Docking 2.4 (HADDOCK 2.4) webserver, using the 8C4L PDB structure of SBSIP and the 6GEH PDB structure of SfSIP and incorporating the substrates Fe(III)-bisucaberin, Fe(III)-alcaligin, NADH and NADPH, in separate runs (56, 57). In each run, 10,000 rigid-body solutions were generated via energy minimization. Subsequently, the 400 structures with the lowest Ambiguous Interaction Restraints (AIRs) underwent semi-flexible simulated annealing in torsion angle space, followed by a final refinement in explicit water. The resulting water-refined structures were clustered using a 1.5 Å backbone Root Mean Square Deviation (RMSD) cut-off and ranked based on their HADDOCK score. Structures with the lowest HADDOCK scores were chosen for further analysis and visualized using PyMOL (31).

## Acknowledgements

The authors are grateful to Anaísa Coelho for teaching the SLIC methodology, to Maria Firmino for providing the sequences for *sip* and *fsr* genes of *S. putrefaciens* DSM 9451, to Filipe Rollo for the assistance with the crystallization robot, to Masaki Fujita for kindly providing the Fe(III)-siderophores putrebactin and bisucaberin, and to Filipe Folgosa for sharing the oxygen-scavenging recipe. The X-ray diffraction data collections were performed at XALOC beamline at ALBA Synchrotron with the collaboration of ALBA staff. This work benefited from access to CERMAX, ITQB-NOVA, Oeiras, Portugal with equipment funded by FCT, project AAC 01/SAICT/2016. Financial support was provided by European EC Horizon2020 TIMB3 (Project 810856). Financial support was also provided by Project MOSTMICRO-ITQB with refs UIDB/04612/2020 and UIDP/04612/2020 and LS4FUTURE Associated Laboratory (LA/P/0087/2020). N-terminal sequencing service was provided by the ITQB Research facilities. Fundação para a Ciência e a Tecnologia (FCT) Portugal is also acknowledged for funding through FCT PT-NMR PhD Program via PD/BD/135187/2017 to IBT.

