## Supplementary material for "Flavin-containing siderophore-interacting protein of *Shewanella putrefaciens* DSM 9451 reveals substrate specificity in ferric-siderophore reduction": suplementary material

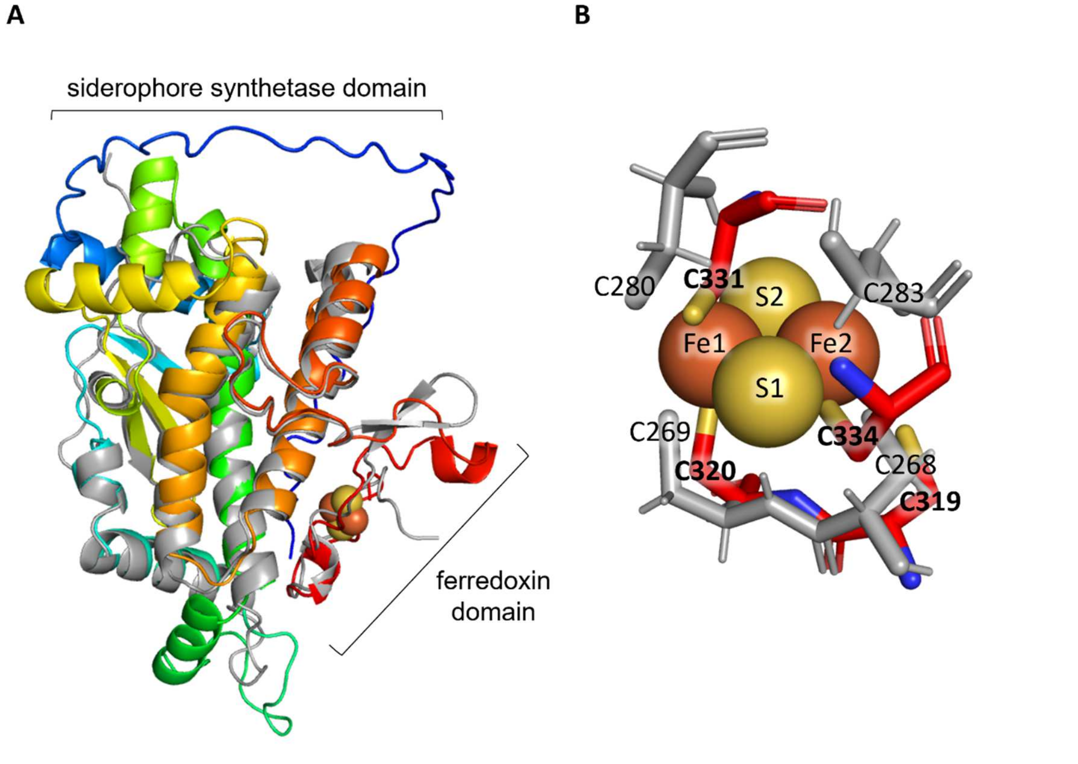


**Fig. S1** Structural comparison of *Sb*FSR with FhuF from *E. coli* **A)** *Sb*FSR structural model (blue to red from the N to the C terminal, AlphaFold model) vs the structure of FhuF (grey, PDB 7QP5) **B)** Close-up to the 2Fe-2S within the ferredoxin domain highlighting coordinating cysteines.

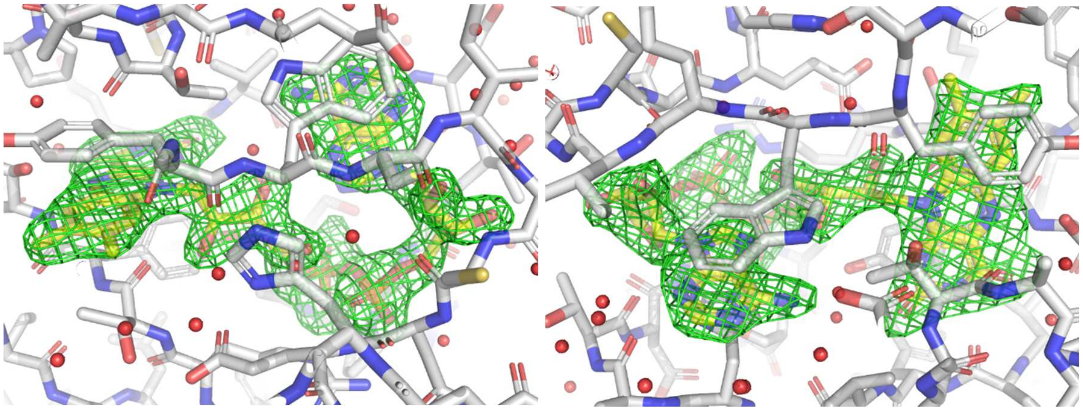

**Fig. S2** FAD omit map for molecule A (left) and B (right) of the crystallographic unit cell

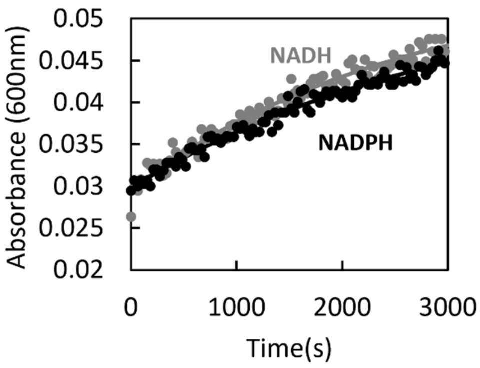


**Fig. S3** Kinetic trace at 600nm of the changes observed after mixing *Sb*SIP with NADH and NADPH. Data for NADH is reported in gray and for NADPH in black.

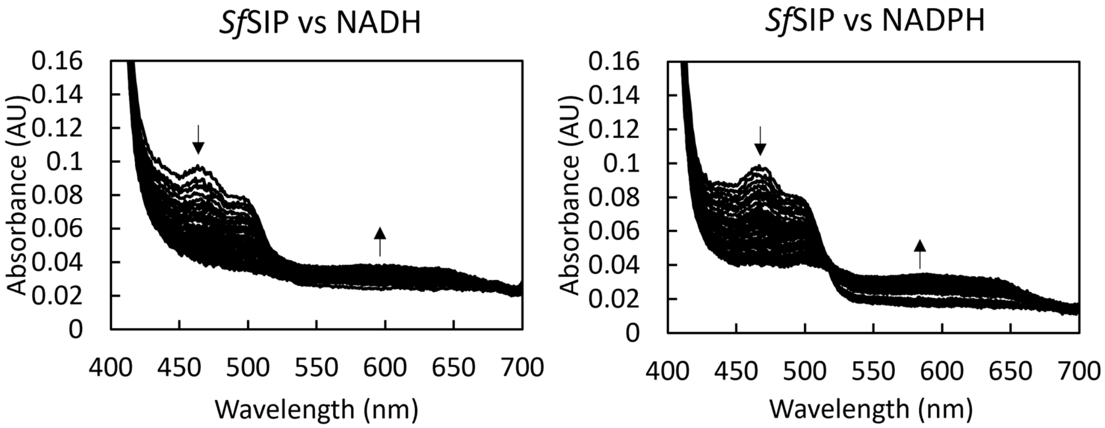


**Fig. S4** Reduction of *Sf*SIP using NADH and NADPH in the presence of an oxygen-scavenging system. **A)** Representative UV-visible spectral changes upon mixing SfSIP with NADH and **B)** NADPH. Arrows indicate the decrease at 470 nm and increase at 600 nm (formation of semiquinone state).

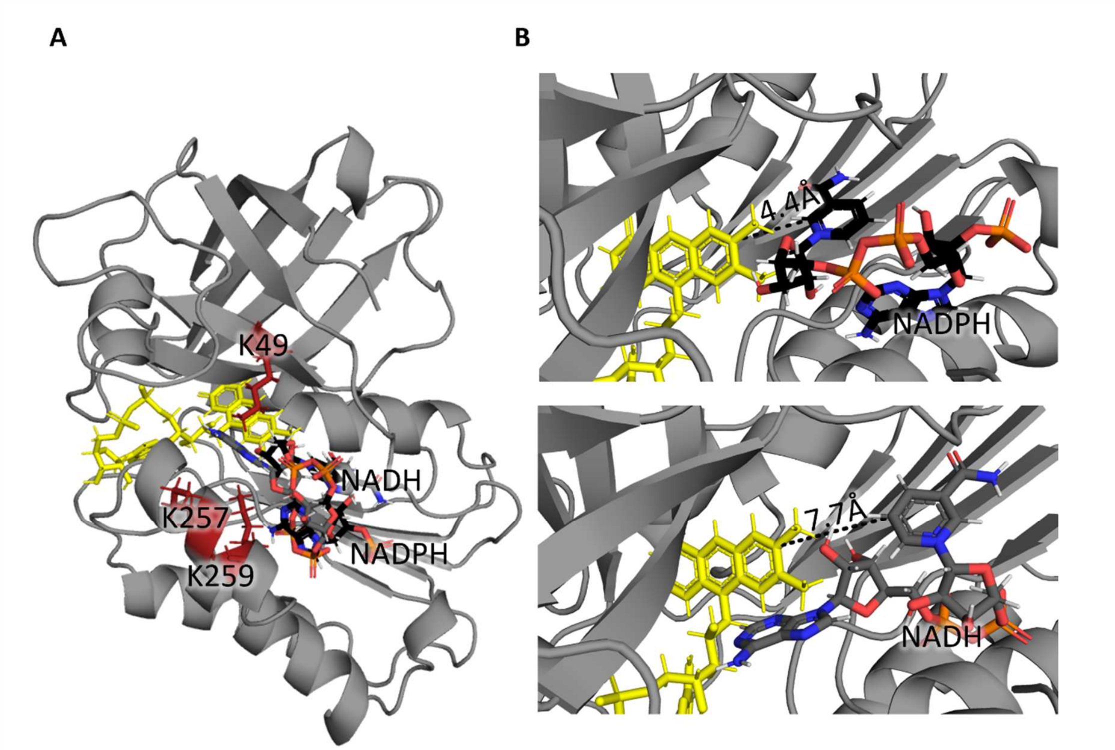


**Fig. S5** Representation of the binding conformations of NADPH and NADH to *Sf*SIP calculated using HADDOCK. A) The binding of NADH (gray) and NADPH (black) occurs in the same region of the isoalloxazine ring of the FAD cofactor (yellow) which is surrounded by the lysine triad (red). (B) zoom of NADPH (top) and NADH (bottom) binding pockets highlighting the shortest distances found between the FAD cofactor and NADPH and NADH respectively.

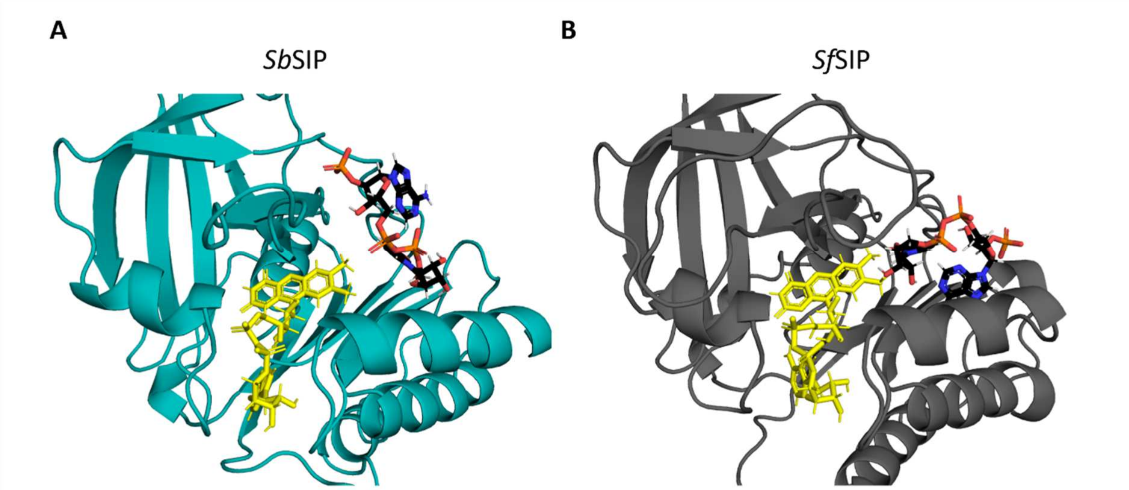


**Fig. S6** Comparison of NADPH (black) binding pocket in *Sb*SIP (teal) (A) and *Sf*SIP (gray) (B) calculated using HADDOCK.

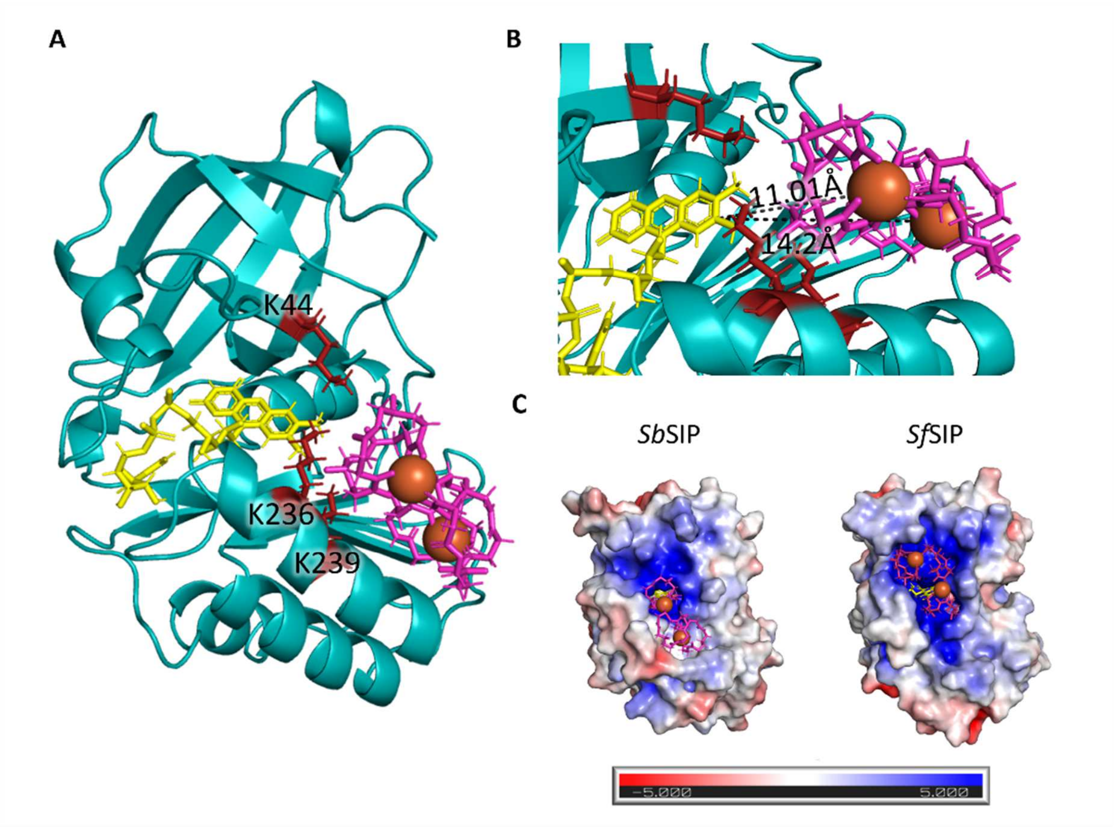


**Fig. S7** Representation of the binding conformations of Fe(III)-alcaligin calculated using HADDOCK: **A)** Docking of Fe(III)-alcaligin (pink) with *Sb*SIP highlighting the lysine triad (red); **B)** Zoom of Fe(III)-alcaligin binding pocket highlighting the shortest distances between the isoalloxazine ring of the FAD cofactor and the two Fe(III) atoms of the Fe(III)-alcaligin complex; **C)** Electrostatic surface potential (−5 to +5 kT/e) of *Sb*SIP and *Sf*SIP and respective Fe(III)-alcaligin binding pockets.

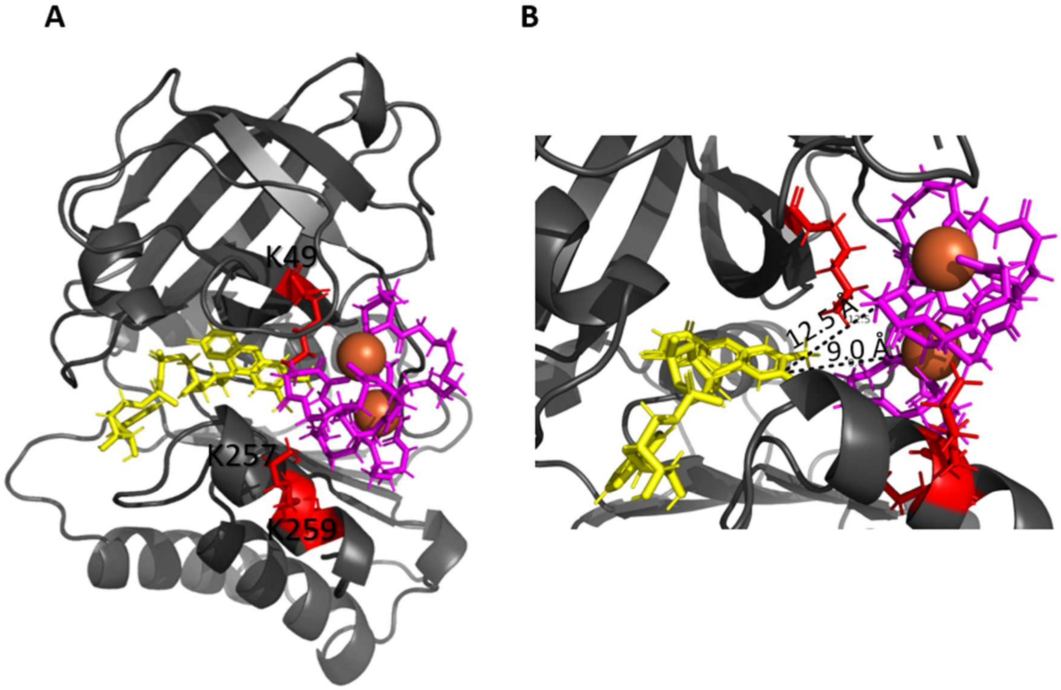


**Fig. S8**. Representation of the binding conformations of Fe(III)-bisucaberin with *Sf*SIP calculated using HADDOCK: A) Docking of Fe(III)-bisucaberin (pink) with *Sf*SIP highlighting the lysine triad (red); B) Zoom of Fe(III)-bisucaberin binding pocket highlighting the shortest distances between the isoalloxazine ring of the FAD cofactor and the two Fe(III) atoms of the Fe(III)-bisucaberin complex.

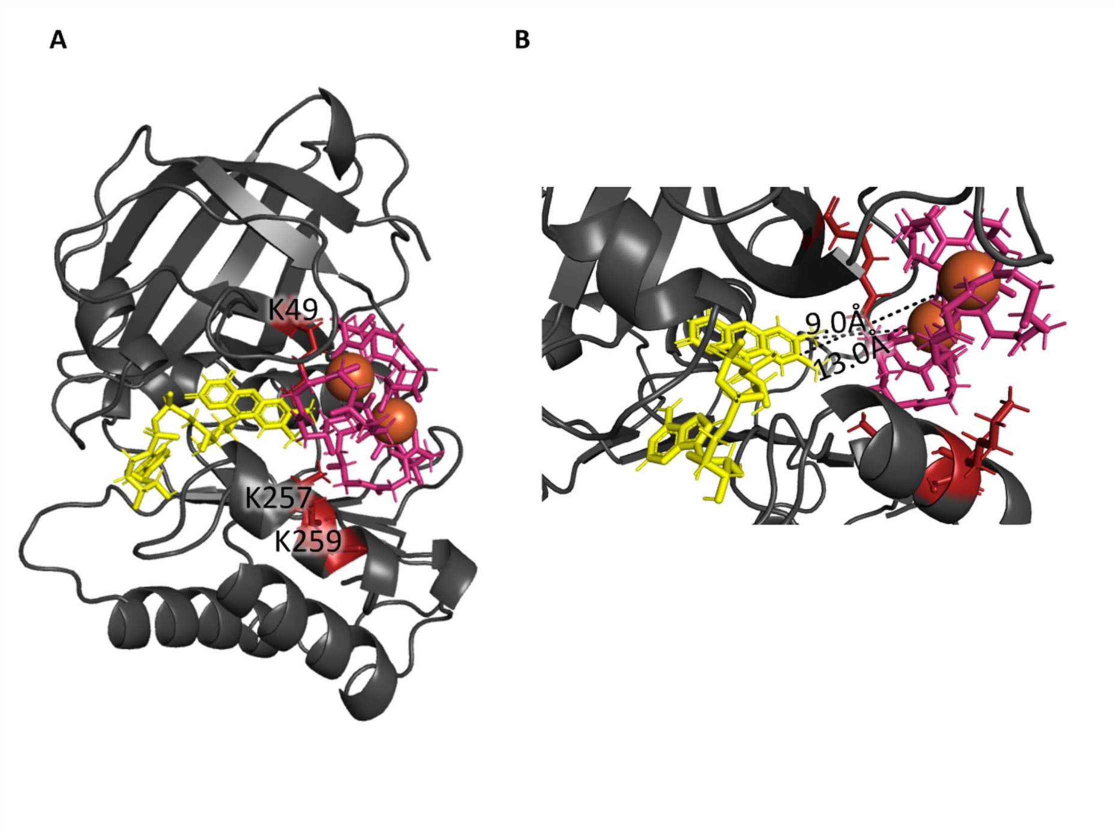


**Fig. S9**. Representation of the binding conformations of Fe(III)-alcaligin with *Sf*SIP calculated using HADDOCK: A) Docking of Fe(III)-alcaligin (pink) with *Sf*SIP highlighting the lysine triad (red); B) Zoom of Fe(III)-alcaligin binding pocket highlighting the shortest distances between the isoalloxazine ring of the FAD cofactor and the two Fe(III) atoms of the Fe(III)-alcaligin complex.
